# *In vivo* two-photon imaging and parasympathetic neuromodulation of pancreatic microvascular dynamics in rats

**DOI:** 10.1101/2020.10.26.355669

**Authors:** Joseph S. Canzano, Narayan Subramanian, Rebeca Castro, Abdurahman Siddiqi, Karim G. Oweiss

**Author notes:** **Correspondence:** Karim Oweiss.

## Abstract

The pancreas has long been known to be densely innervated with parasympathetic, sympathetic, and visceral afferent fibers that are believed to exert significant influence on local endocrine activity and vascular function. Yet the extent to which these interactions depend on neurovascular dynamics in the normal and pathological states remain largely unknown. Herein we describe a new method for high resolution functional imaging of the rat pancreas *in vivo*. The method comprises a number of elements: a stability-optimized preparation in dorsal recumbency immobilizing several square centimeters of intact pancreas for upright fluorescent imaging while leaving access for concurrent manipulation of abdominal nerves, a full-frame two-photon imaging protocol and analysis pipeline supporting high-throughput (100+) monitoring of islet and acinar microvessel diameter dynamics simultaneously, and a first adaptation of random-access linescan imaging to the pancreas capable of tracking internal blood flow speeds up to 5 mm/s at 20 Hz across multiple microvessels. These methods were then deployed in concert to characterize the capacity of parasympathetic fibers to modulate pancreatic microvascular dynamics with compartment specificity. Electrical stimulation was repeatedly applied to the abdominal vagal trunks at various current magnitudes while imaging islet and acinar microvascular populations in the pancreas. Vagal stimulation consistently elicited increases in both islet and acinar capillary population motility in a current-dependent manner, with only acinar responsive vessels trending toward dilation. Further, we found vagal stimulation to profoundly and reversibly disrupt all traces of fast-wave vasomotor oscillation across a lobular arteriole-venule pair, and this was associated with a significant increase in average flow speed. Together, these findings add to mounting evidence that vagal projections exert tangible reversible influence on pancreatic microvascular activity and underscore the potential for new neuromodulation-based strategies to address diabetes, pancreatitis, or other diseases of the pancreas under autonomic nervous influence.

## Introduction

The pancreatic Islet of Langerhans is a highly complex signaling environment. Comprised of neuroendocrine cells, projections from all arms of the peripheral nervous system, and the various cellular constituents in a highly specialized capillary plexus, competing influences ultimately culminate in secretory activity to precisely control blood glucose. While direct interactions between peripheral axons and β or α cells through non-classical synapses have been described in rodents, no analogous structural connectivity has yet been demonstrated in humans ^1–3^. Contemporary theory therefore suggests that nervous control of islet secretion principally occurs indirectly through modulating islet blood flow via contractile activity among islet pericytes and arteriolar smooth muscle cells ^4–6^. Neurovascular studies are therefore of utmost importance to uncover the basic principles of neuro-insular dynamics, and to develop neuromodulation-based therapies to address diseases of the islet such as diabetes.

Unfortunately, these studies are not commonplace due to the exceptional difficulties associated with the thorough examination of islet neurovascular interactions in their natural microenvironment. In particular, the pancreas is a soft abdominal organ filled with digestive fluid, subjected to movements of the diaphragm, and mechanically bonded via connective and vascular tissue to neighboring organs including spleen, stomach, duodenum, intestine, and liver. Consequently, most functional studies of pancreatic microvascular or related neural processes are either carried out *in vitro*^7^, *ex situ* ^8^, or *ex vivo* ^9,10^. In limited cases *in vivo* pancreatic imaging has been described in mice, but this has been done in a setup where limited access to abdominal nervous structures was available, and the compatibility of these approaches have not been demonstrated in studies involving autonomic control or neurovascular activity ^11–13^.

Herein we set out to develop a surgical preparation for upright, intravital imaging of pancreatic microvascular dynamics in rats with sub-capillary resolution, tailor-made for neurovascular studies by providing access to surrounding abdominal structures. Two-photon laser scanning microscopy (TPLSM) was chosen as the ideal imaging modality for its high resolution and superior signal to noise ratio *in vivo* ^14^, and a series of experiments were then performed to verify that these combined techniques enable thorough characterization of various pancreatic microvascular responses to parasympathetic activation. The abdominal vagal trunks were chosen as a parasympathetic target as their connectivity to and within the pancreas is well described in rodents ^1,4,15^, the abdominal locus is expected to avoid confounding effects on cardiac and respiratory activity ^16^, and vagal stimulation is currently undergoing clinical development for humans as a treatment for obesity ^17^, making its potential impact on understanding digestive organ physiology highly valuable. By simultaneously examining diameter changes among acinar and islet capillary populations, as well as internal flow speed dynamics across terminal microvessels, we found numerous distinct and unexpected functional responses to vagal stimulation. We believe that the detailed methods and analytical tools provided herein will spark greater interest and further guidance to the study of pancreatic function *in vivo*.

## Results

### Two-Photon Imaging of Pancreatic Microvessel Networks at Sub-Capillary Resolution

The functional requirements for the preparation and imaging parameters were first determined in order to meet the described goals. First, as pancreatic capillaries can be as small as 4 μm in diameter, the sample must be stabilized on the scale of nearly single microns to avoid uncorrectable movements in the axial (z) direction. To minimize impact to normal vascular physiology, the pancreas must be exteriorized and mechanically stabilized without any compression, stretching, drying, or cooling, and maintained at the same elevation relative to the heart, all while attached to an anesthetized animal in dorsal recumbency to leave access to abdominal nerves. Targeting of individual nerves or catheterization of small blood vessels requires the use of a surgical microscope, so ideally sensitive surgical steps would be performed outside the physical constraints of the typical upright two-photon imaging stage before imaging.

The developed procedure meeting these requirements is illustrated in Figure 1A-B. The animal lies on a height-adjustable stage built on a 12” × 12” optical breadboard to allow the mounting of clamps for tubing and imaging pedestal, while being easily movable between surgical and imaging stations. Abdominal access is achieved through a T-shaped incision along the linea alba, and the pancreas along with spleen are sparsely adhered to a temperature-controlled aluminum imaging pedestal inserted to roughly maintain their original positions. All major blood vessels traversing the pancreas and spleen are left intact and only connective tissues are dissected (i.e. the gastrosplenic ligature). As the pancreas shares many immutable mechanical linkages to the stomach and liver it became necessary to decouple respiratory movements with the anterior abdomen to meet stability goals. A bilateral pneumothorax is therefore created by a single crosswise incision through the diaphragm, and breathing is subsequently maintained via tracheostomy and artificial ventilation. Carefully monitored ventilated animals remain stable for many hours, and due to the improved displacement of CO_2_ from the lungs, diaphragm contractions associated with inspiratory attempts are eliminated without the need for respiratory paralytics which could further impact vascular physiology. In addition to the information included in the online methods section of this manuscript, a detailed step-by-step protocol for this procedure has been made freely available on protocols.io ([link](https://www.protocols.io/view/in-vivo-two-photon-imaging-and-parasympathetic-neu-bm3tk8nn)).

**Figure 1.**
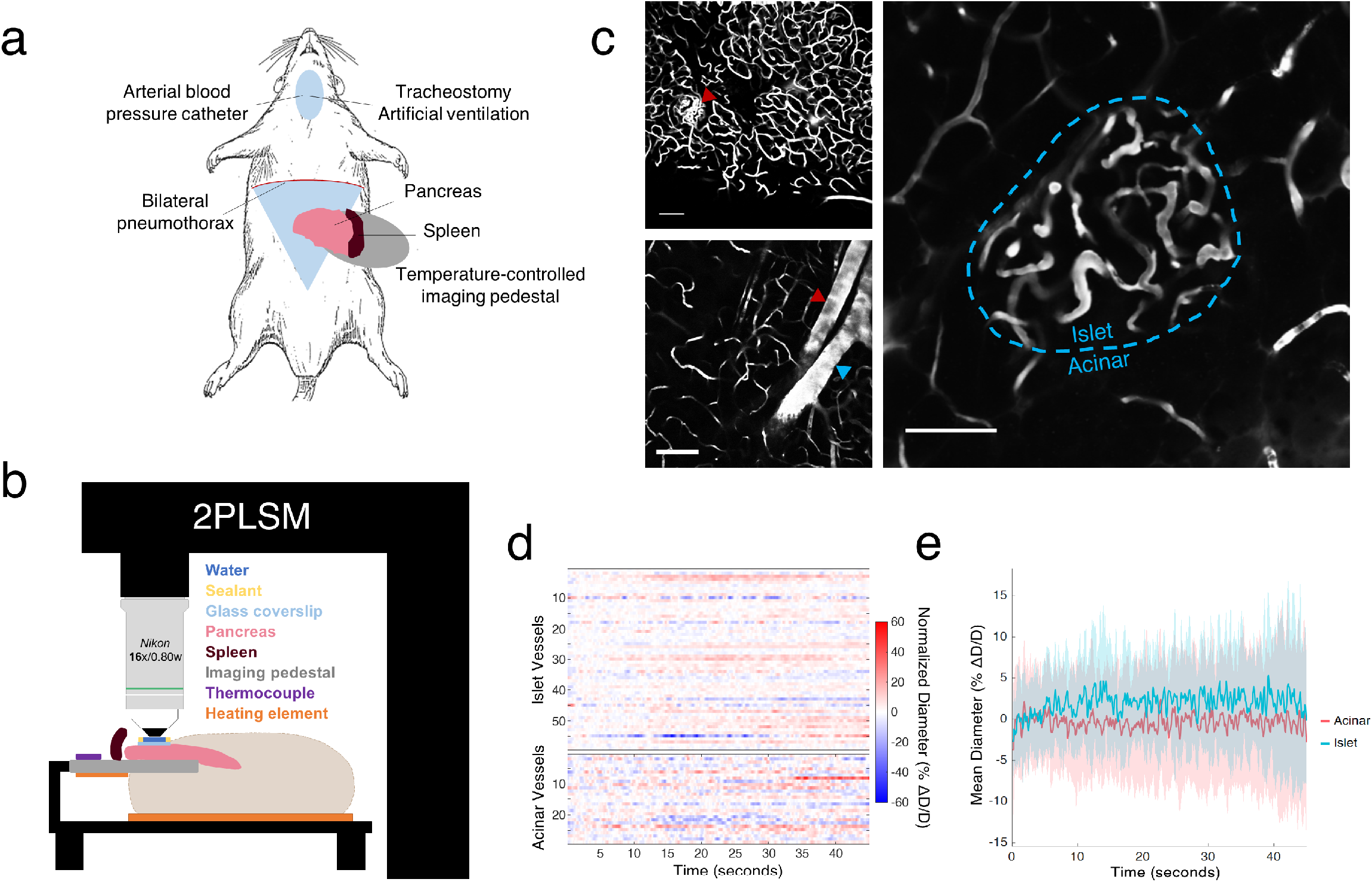
Overview of surgical preparation and full-frame *in vivo* imaging procedure. A) Main features of surgical preparation. B) Layout of pancreas mounting and optical access to the sample. The pancreas and spleen are sparsely adhered to a heated aluminum imaging pedestal positioned to roughly maintain their original elevation and position. A small round coverslip is placed directly atop a region of pancreatic tail, and a water interface for imaging is created by sealing the coverslip perimeter with Vaseline. All exteriorized abdominal surfaces are covered in soaked gauze and kept moist to prevent fluid loss. C) Appearance of pancreatic microvasculature after i.v. injection of dextran-FITC. Top left, z-projection (40 μm) of showing microvascular network from a pancreatic lobule. Islet at red triangle. Bottom left, representative acinar region featuring capillaries and lobular arteriole-venule pair at red and blue triangles, respectively. Right, representative FOV featuring both islet and peri-islet acinar capillaries with border shown in blue. Islets are easily distinguished by their distinct glomerular microvascular morphology. Scalebars: 100μm. D) Example raster plot of islet and acinar capillary diameters recorded simultaneously, measured from baseline recording. E) Islet and acinar capillary diameter activity are analyzed as separate populations. Baseline mean +/- SEM from example recording.

If stabilization and mounting are carried out carefully, this procedure provides upright imaging access to several cm^2^ of pancreatic tail with near sub-micron mechanical stability for several hours. After i.v. injection of a fluorescent tracer such as dextran-FITC, whole microcapillary networks can be visualized via TPLSM with superb contrast from parenchyma, including clearly defined erythrocyte shadows flowing within each microvessel (Figure 1C, Supplemental Video 1). At 1x optical zoom through a 16x Nikon water-immersion objective providing 800 μm^2^ FOV, the microvascular network of near whole pancreatic lobules can be imaged simultaneously at subcapillary resolution at depths approaching 100 μm before scattering becomes too significant to resolve most capillaries (Figure S1^a^). Superficial islets are readily located and identified without difficulty by the unique glomerular characteristics of their capillary beds, and the FOV range at the standard 16x allows large numbers of islet and acinar microvessels to be imaged simultaneously and analyzed as separate populations (Figure 1D,E). In the acinar space, lobular arterioles and venules can be found of all sizes, isolated or traversing as pairs (Figure 1C). We also found that fast-wave vasomotor oscillations, well known to be exhibited by arterioles^18^, can be readily visualized as a rhythmic, pulsatile narrowing in diameter among these vessels (discussed in greater detail in the following sections).

### Stabilized Mounting Facilitates Random-Access Linescan Imaging to Monitor High-Speed Flow Dynamics

Visualizing the flow of erythrocyte shadows makes quantification of internal flow speeds possible, but it became immediately apparent that many pancreatic microvessels exhibit internal flow speeds exceeding what can be resolved by full-frame resonant scans. Notably, this included many islet capillaries and all arterioles, which would be the primary actuators of interest if the goal is to devise a neuromodulation-based control approach over compartment-specific microvascular activity. Thus, random-access linescan imaging ^19^ was adapted within this preparation to compliment the high-throughput full-frame diameter measurements with a temporally-advantaged counterpart. In combination with our stability-optimized protocol, this approach readily achieves internal flow speed measurements from multiple pancreatic microvessels up to 5 mm/s at sampling rates exceeding 20 Hz (Figure 2A-D). Velocity recordings usually have complex time-varying profiles, including prominent oscillations at frequencies attributable to both cardiac pulsation and vasomotor oscillation (Figure 2C-F). While not all pancreatic microvessels sized 20-100 μm exhibited diameter oscillations, flow speed oscillations in the vasomotor frequency range (~0.1-0.6 Hz) were almost always found in linescan recordings, even within capillaries (Figure S2). Finally, as this imaging modality supports the simultaneous measurement of microvessel diameter and internal flow speed, correlation and phasic analysis can be performed between the two variables within and across vessels, though phase ambiguity can be troublesome to resolve for vessels possessing highly-regular oscillations, in the absence of any disruptive perturbations (Figure 2E).

**Figure 2.**
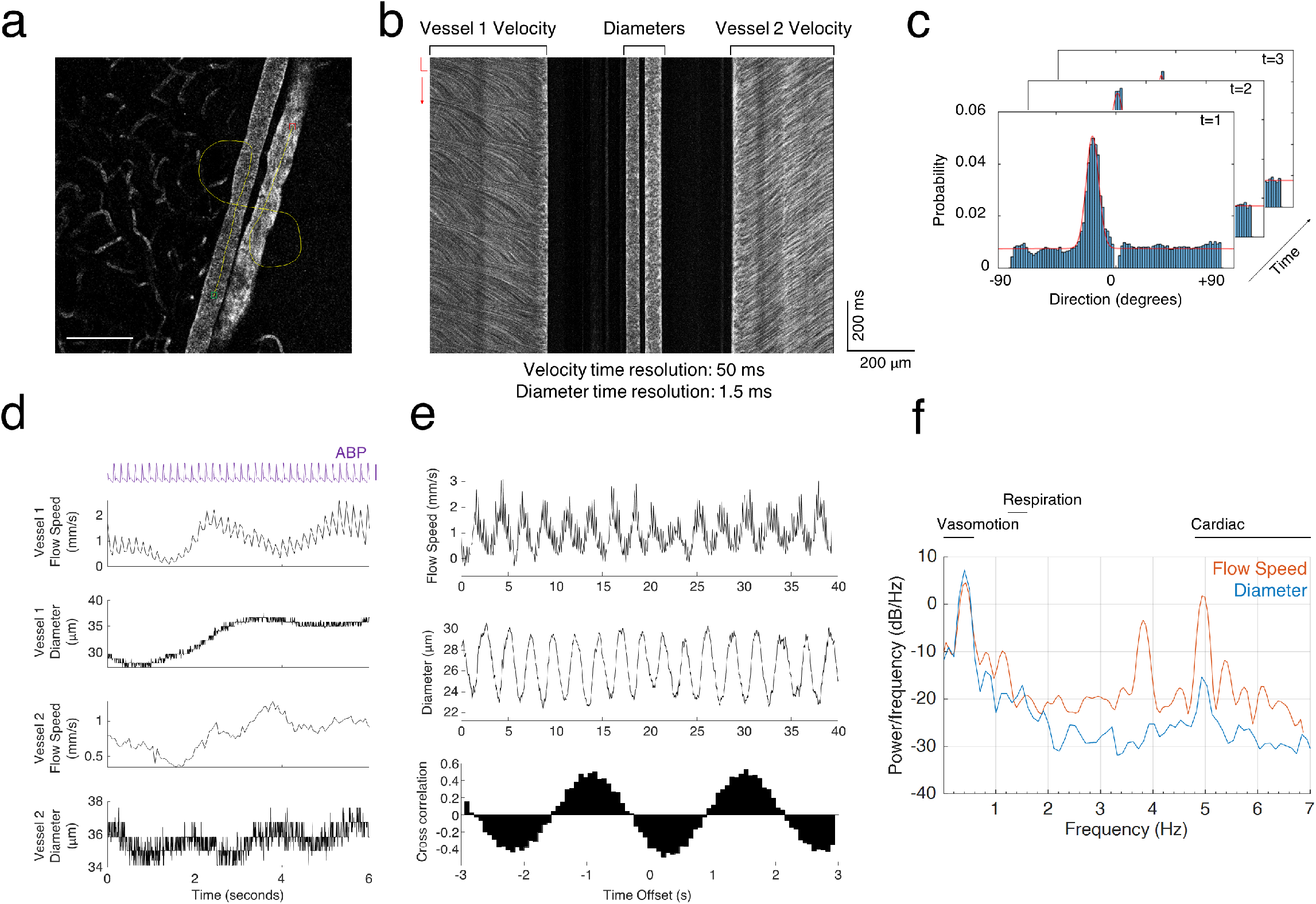
Overview of linescan imaging and recording features. A) The imaging laser is directed along a freehand path, both along and across microvessels of interest, to respectively capture flow speed and diameter dynamics. The yellow line (start at green, end at red) depicts an example path imaging an arteriole-venule pair. Scalebar, 100 μm. B) Appearance of the resulting linescan collection with columns labeled for their subsequent analysis. Erythrocyte shadows traversing the vessel appear as dark lines with slopes proportional to distance/time, so velocity columns are first windowed (red bracket) and the slope content within each becomes one velocity timepoint. C) Intermediate outputs from the slope detection step. The directional content of each velocity time bin is estimated (see methods), and the resulting histograms are gaussian fit to obtain the peak direction, which is geometrically related to distance/time. D) Resulting flow speed and diameter traces from the vessels in A-B, along with concurrent arterial blood pressure recording. Putative arterioles (vessel 1) can be identified at this stage by the strong presence of cardiac oscillations in the flow speed along with high degree of diameter flexibility-note the lack of both in vessel 2. ABP scalebar, 50 mmHg. E) Top two, extended baseline linescan recording data from another representative putative arteriole possessing clear vasomotor oscillations in both diameter and flow speed activity. Bottom, cross-correlation can be calculated for flow speed vs diameter showing the phasic relationship between the two. F) Power spectral density estimate for the representative arteriole in E, showing prominent peaks in vasomotor band (slow and fast-wave included, 1-25 cpm). Note the presence of strong cardiac oscillations (275-400 bpm) in the flow speed and lack of respiration (75-100 bpm, locked by ventilator) artifacts.

### Vagus Stimulation Elicits Distinct Shifts in Islet and Acinar Capillary Network Diameter Dynamics

A series of experiments were performed to demonstrate the power of these techniques and provide a first look at how parasympathetic activation may influence pancreatic microvascular dynamics between compartments. Repeated electrical stimulation was delivered to the abdominal vagus nerves with varied current amplitude while imaging the pancreatic microvasculature using the approaches described above: either full-frame imaging (1024×1024, 15 Hz) to record diameter from a large amount of microvessels simultaneously, or fast linescans (0.8-1.2 KHz) to record internal flow speed and diameter from select microvessels (Figure 3A). A total of 89 full-frame recordings were collected from n=2 adult rats, 60 of which including both islet and acinar capillary beds, averaging 32.3 acinar and 43.1 islet microvessels per field-of-view (Figure 3B).

**Figure 3.**
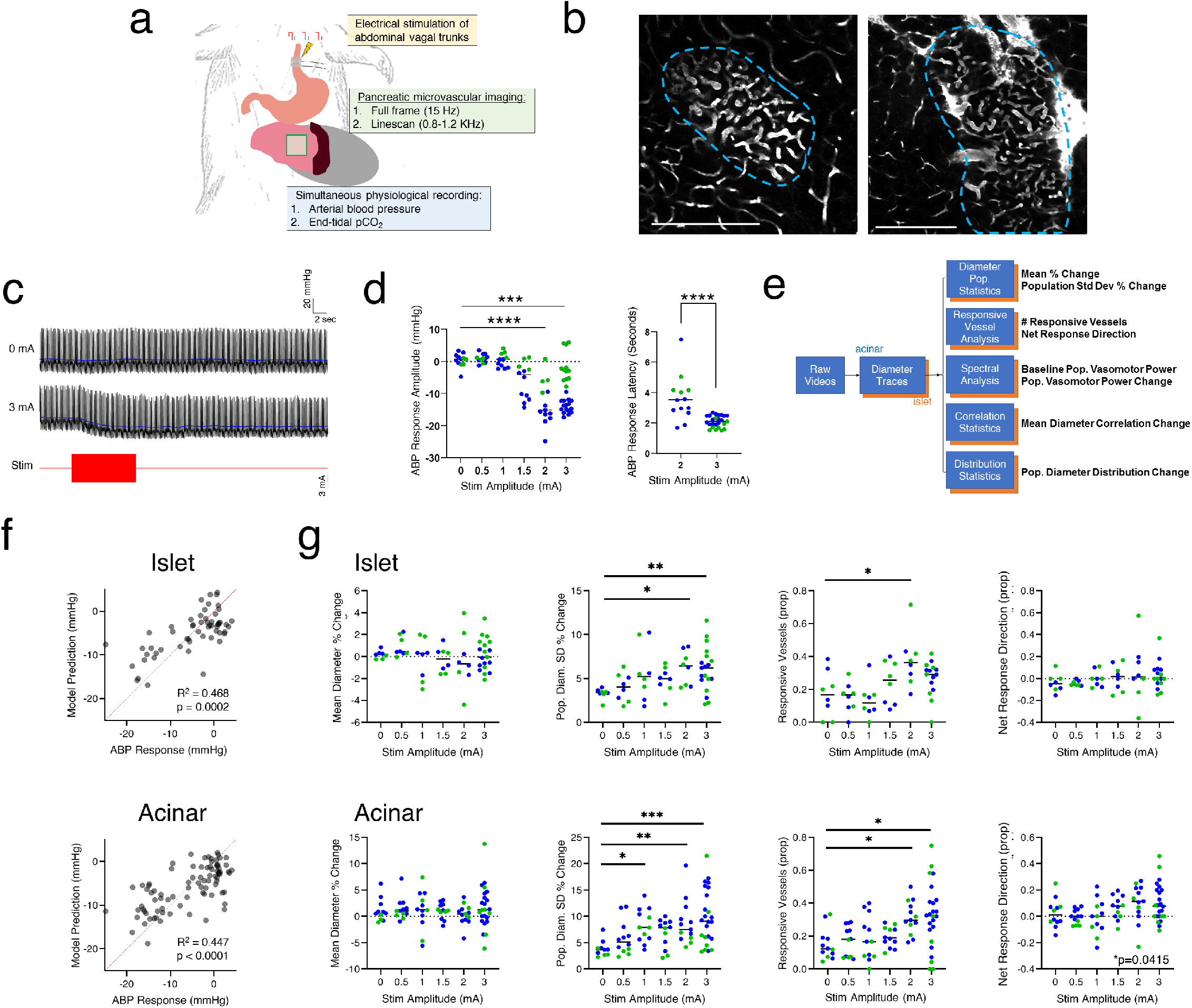
Abdominal vagus stimulation impacts microvessel diameter dynamics in both islet and acinar compartments. A) Schematic of experiment. The basic procedure in Fig 1A-B is appended with a bipolar cuff electrode implanted on the esophagus to activate both abdominal vagal trunks with biphasic pulse trains (800 μs, 50 Hz, 10 second duration) of varied current amplitude (0-3 mA) in repeated, randomized trials. B) Example islet + acinar FOV’s analyzed with concurrent stimulation. Islet/acinar border in blue. Scalebar, 200 μm. C) Example arterial blood pressure (ABP) waveforms recorded during control (0 mA) and stimulation (3 mA) trials showing drop in systemic blood pressure associated with abdominal vasodilation following stimulation onset. Blue lines show lowpass-filtered waveform used to determine response amplitude (see methods). D) Left, current amplitudes of 2 mA and above elicited a significant transient drop in ABP following stimulation onset (2mA: n=13, p<0.0001; 3mA: n=28, p=0.0007). Right, Above the ABP response threshold, higher stimulation current led to shorter response latency (Mann-Whitney p<0.0001). E) Overview of processing pipeline and main analysis outputs for full-frame data. Islet and acinar microvessels are analyzed as separate populations and outputs are relative to a five second baseline period included in each recording. F) The microvascular dynamic variables in E can predict the degree of vagal activation indicated by the ABP response, using either acinar or islet microvessel data (p-values are ANOVA versus constant model; stimulation amplitude was also fit with similar results, Supplemental Figure 4). G) Select microvascular population statistics plotted against stimulation amplitude for islet and acinar regions separately. While mean population diameter was unaffected by stimulation (column 1), the variability of diameters (SD % change vs baseline) is significantly increased by high current amplitudes for both islet and acinar capillaries compared to 0 mA trials (column 2). The proportion of vessels that significantly deviate from baseline is also increased among islet and acinar vessels at high current amplitudes (column 3), with only acinar responsive microvessels trending toward vasodilation under these conditions (column 4, acinar Kruskal-Wallis p=0.0415, however no MC-corrected comparisons reach significance). Plots of the remaining outputs from E are shown in Supplemental Figure 5. All points represent single recordings and are colored by experiment where applicable. Horizontal bars denote group median.

Electrical stimulation of the abdominal vagus consistently elicited a significant transient drop in arterial blood pressure for current amplitudes at or above 2 mA (Figure 3D, 2mA p<0.0001, 3mA p=0.0007 compared to 0 mA control group). Within this range, higher current led to a shorter vasodilation response latency (Figure 3D, p<0.0001). Additionally, no significant change from baseline in pulse rate, pulse pressure, or peak expired pCO2 were found as a result of stimulation (Figure S3). As vasodilation of abdominal blood vessels would be expected to decrease the systemic blood pressure (more vessel area for same fluid volume), these results indicate that the abdominal vagi were activated as intended, and targeting the abdominal trunks succeeded in avoiding significant confounding cardiac or respiratory effects.

We then sought to examine the full-frame recordings for any effects of vagal stimulation on islet and/or acinar microvascular dynamics. As registering individual microvessel segments between trials is difficult (see discussion) an analysis was designed focusing on metrics that do not require repeated samples from the same population of vessel segments but could still cover any changes in time-varying diameter activity one might expect. An overview of the resulting statistical outputs is provided in Figure 3E featuring the following metrics versus baseline values where applicable: changes in basic diameter population statistics (population mean and SD), the proportion and net movement direction of “responsive vessels” (defined as those which diameter crossed +/- 4 baseline standard deviations during stimulation period), changes in fast-wave vasomotor oscillation power (frequencies 0.3 - 0.6 Hz), changes in the mean correlation between all vessels (grouped by islet or acinar), and several common metrics capturing shifts in diameter distributions. All metrics were designed and implemented prior to any data analysis. The full analysis pipeline including usage documentation has also been made freely available on protocols.io ([link](https://www.protocols.io/view/in-vivo-two-photon-imaging-and-parasympathetic-neu-bm3tk8nn)).

As a first exploration of whether meaningful trends exist in the data, the aforementioned microvascular variables were used to fit generalized linear models predicting either the stimulation current applied or the magnitude of the vasodilation response from the ABP recordings. Both of these models were found to have significantly more predictive power compared to a constant model using either islet (ABP p=0.0002, Stim Amp p=0.0032 vs constant model) or acinar (ABP p<0.0001, Stim Amp p=0.0028) variables, indicating the presence of causal influence of vagal stimulation on microvessel dynamics in both pancreatic compartments (Figure 3F, Figure S4). As some dynamic variables were correlated, this process was repeated after performing principal components analysis to identify decorrelated latent structures in the data most critical for model performance, the results of which are discussed further in Figure S4.

Examining the individual variables led to greater insight into the specific microvascular dynamics impacted by vagal stimulation (Figure 3G). No significant shift in mean population diameter was found in response to any current amplitudes tested compared to 0 mA control trials (Fig 3G, column 1). However, a significant increase in diameter variability was observed over control trials, in both islet and acinar compartments, with the effect size scaling with current magnitude (Fig 3G, column 2; islet 3mA p=0.007, acinar 3mA p=0.0004). These results indicate that while vagus stimulation induced an increase in microvessel motility, the net shift in the whole population was not coherent and large enough to significantly change the mean diameter, a somewhat expected result as a notable feature of recordings is the high variability of responses between individual capillaries. As averaging can be lossy, it could be reasoned that a significant and coherent shift among a subset of “responsive” capillaries may still be present in the data and merely obscured in the mean by those which do not respond to stimulation. In addition, vessels within the same compartment could be responding in separate directions based on some unseen subtype distinction leading to a net zero change. Ultimately, we indeed found a greater proportion of responsive vessels at high stimulation currents compared to low currents in both islet (p=0.0500) and acinar capillaries (p=0.0106; Figure 3G, column 3). Interestingly, the net shift direction among responsive vessels was toward dilation in only acinar capillary beds (Kruskal-Wallis p=0.0415) whereas islet responsive capillaries showed no directional bias (Kruskal-Wallis p=0.447; Figure 3G, column 4). No stimulation currents tested led to a significant change in the average correlation between capillaries, global vasomotor oscillation power, or the overall distribution of diameters among islet or acinar microvascular networks compared to control trials (Figure S5; one distribution metric approached significance for acinar). Altogether these results suggest vagal stimulation elicits a significant shift in select diameter dynamics in both islet and acinar microvessel populations, best characterized as a global increase in capillary motility, but notably a greater degree of vasodilatory coherence in the acinar response compared to islets.

### Vagus Stimulation Transiently Disrupts Diameter and Flow Traces of Fast-Wave Vasomotion

We also examined microvessel diameter and flow speed signatures of vasomotor oscillations for dynamic changes associated with vagus stimulation. Several oscillating microvessels were encountered over the course of this study^b^ (Figure 4A, Supplemental Video 2) and in one representative lobular arteriole-venule pair, both full-frame and linescan recordings were collected over repeated trials stimulating the abdominal vagus. Strikingly, vasomotor diameter oscillations exhibited by the putative arteriole could be consistently and completely eliminated by vagal stimulation (Figure 4B,C). All current magnitudes at or above 1.5 mA were associated with a significant transient decrease in diameter oscillation power compared to 0 mA control trials (Figure 4D, horizontal bars denote p<0.05 vs 0 mA), effectively increasing the average diameter during stimulation. Notably, the latency for this vasomotor disruption was consistent with that observed in the arterial blood pressure response to vagal stimulation discussed earlier (Figure 3D).

**Figure 4.**
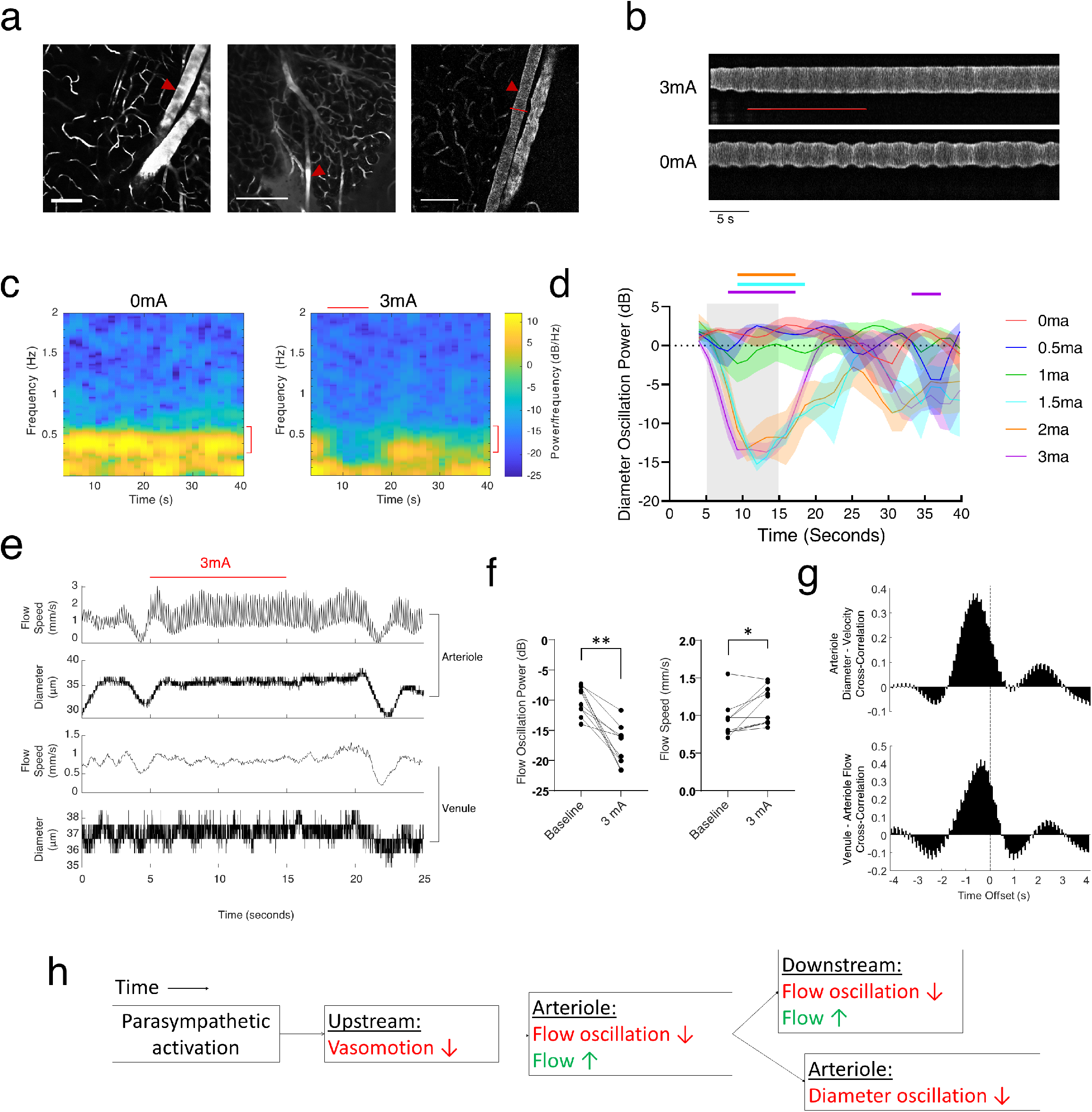
Vagus stimulation can disrupt traces of microvascular vasomotor oscillation in both diameter and velocity activity. A) Example microvessels with strong diameter oscillations (red triangles) encountered in different animals. The lobular arteriole-venule pair in the rightmost FOV were analyzed as representative vasomotor vessels in a vagus stimulation experiment (Fig 3A). B) Diameter oscillations exhibited by the arteriole were almost completely disrupted by vagus stimulation (line profile taken from red bar in A; horizontal red line is stimulation period). C) Average spectrograms computed from repeated trials delivering 0 mA (n=4) or 3 mA (n=8, horizontal red line is stimulation period) current amplitude show consistent disruption of diameter oscillations in fast-wave vasomotor band in 3 mA trials (red brackets). D) Diameter vasomotor oscillation power is significantly reduced at current amplitudes >=1.5 mA versus 0 mA trials. Traces are mean +/- SEM from repeated trials. Horizontal colored bars represent timepoints reaching significance criterion (p<0.05, multiple t-tests with Holm-Sidak correction). 0-2mA groups all have n=4 replicates, 3mA n=8 replicates. E) Representative linescan recording from the same vessels (Figure 4A, right). Simultaneous measurement of diameter and flow speed show flow speeds are correlated across both vessels and also transiently disrupted by vagus stimulation (red horizontal line). Note the lack of cardiac oscillations in putative venule flow speed as well as stationary diameter compared to putative arteriole. F) Quantification of flow speed responses to repeated trials of vagal stimulation among both vessels. Left, flow speed oscillations were significantly disrupted during stimulation compared to baseline (paired Wilcoxin p=0.0020, n=10) within both vessels. Right, the disruption of oscillations during stimulation were associated with an increase in average flow speed (Wilcoxin p=0.0137, n=10) within both vessels. G) As these disruptions alleviate phase ambiguity, evidence of causal relationships can be obtained by analyzing delays between traces. Top, cross-correlation (average of n=7 recordings) between arteriole diameter and velocity traces show arteriole diameter lags flow speed by 650 ms. Bottom, venule flow speed lags arteriole flow speed by 250 ms (average of n=7 recordings). Vertical line denotes no delay. H) Vasomotor response to vagus stimulation findings summarized diagrammatically.

Simultaneous internal flow speed and diameter measurements from the same arteriole-venule pair uncovered more vasomotor effects of vagus stimulation (Figure 4E). First, arteriole flow speed oscillations were also found to be consistently disrupted by vagus stimulation (p=0.0020) leading to an increase in average flow speed within the arteriole compared to baseline (Figure 4F; p=0.0137, n=10). As this essentially introduces a perturbation resolving phase ambiguity, multiple predictions can be made regarding the relative timing of events within and across these microvessels (and implicitly, flow through their adjoining capillaries). For example: as expected, paired arteriole and venule flow speeds were clearly correlated, with changes in venule flow lagging arteriole flow by about 250 ms (Figure 4G, bottom). More importantly, arteriole diameter and flow speed were also correlated, with diameter lagging internal flow fluctuations by about 600 ms, suggesting flow oscillations cause diameter oscillations, and not the other way around (Figure 4G, top). These data support the simple model in Figure 4H in which abdominal vagus nerve stimulation drives changes in vasomotor dynamics within these microvessels. An unobserved source of flow oscillations upstream is transiently disrupted by parasympathetic influence leading to an elimination of arteriole flow oscillations, subsequently eliminating its diameter oscillations and affecting velocity oscillations observed downstream. Decreases in flow oscillations are associated with increases in overall flow speed and presumably greater fluid flux, eventually leading parasympathetic activation to increase overall perfusion through this microvascular network (as expected within a digestive organ). As only one representative arteriole-venule pair were used, this model is not expected to globally describe these phenomena among the entire acinar microvasculature. More appropriately, the model suggests that, for some pancreatic microvascular networks, this cascade of events can be reliably evoked by vagus stimulation. Furthermore, it serves to demonstrate the power of the combined techniques in inferring causal links between microvascular dynamics and neural events within pancreatic tissue *in vivo*.

## Discussion

As knowledge of the interplay between peripheral innervation, vascular actuators, and pancreatic secretory activity deepens, great need still exists for methods to functionally interrogate these complex systems *in vivo* without disrupting the native pancreatic microenvironment. The contributions of the current work towards *in vivo*, intact imaging of the pancreas in rats are multifold. First, we described a surgical preparation in dorsal recumbency immobilizing several cm^2^ of intact pancreas for upright imaging while leaving access to sensitive abdominal structures. Second, we developed a full-frame TPLSM imaging protocol and analysis pipeline supporting the simultaneous monitoring of diameter dynamics among islet and acinar microvessel populations at sub-capillary resolution. Third, we adapted random-access linescan imaging to the pancreas achieving simultaneous tracking of intralaminar flow velocity and vessel diameter from multiple microvessels at speeds up to 5 mm/s at 20+ Hz. Fourth, we deployed these methods in concert to functionally characterize compartment-specific pancreatic microvascular responses to electrical stimulation of the abdominal vagus nerve, yielding specific insights into the nature of microvessel action within parasympathetic influence.

Several existing methods to achieve *in vivo* imaging of either transplanted islets in the retina or in their original microenvironment have been described^13^ and elegantly applied in studies of immune cell migration ^12^, architectural features of the islet capillary plexus ^11^ and neurotransmitter modulation of islet pericyte contractility ^5^. While the preparation described here was applied to examine parasympathetic control of pancreatic microvascular dynamics, it is directly transferrable to functional neurovascular studies entirely within the islet, which would also benefit from greater ease targeting abdominal nerves, broad optical access, and a high degree of stability. It remains to be seen the extent to which the full array of modern optical neuroscience tools such as modern genetically-encoded calcium indicators ^20^, engineered opsins for cellular resolution optogenetic stimulation ^21–23^, and/or targeted ablation ^24^ could be fully leveraged in the pancreas *in situ*. Rats, while convenient for interacting with individual branches of peripheral nerves and the complicated celiac plexus, are disadvantaged models for such studies for the limited genetic toolset, but this issue is becoming only merely inconvenient as viral methods for transgene expression improve at a rapid pace ^25^. Most noteworthy is the onset of modern engineered adeno-associated virus (AAV) serotypes capable of widespread peripheral infection with one intravenous injection ^26^, which would be expected to show dramatic improvements in efficiency compared to older-generation AAV tools that were already sufficient to express optical labels in pancreatic tissue and even beta-cells *in vivo* ^27–29^.

While the effects of selective denervation and/or application of neurotransmitters on islet secretion and perfusion have been well-covered in studies dating back several decades ^1,2,6,30,31^, the potential for peripheral neuromodulation to selectively drive pancreatic activity remains purely theoretical to our knowledge. At the outset of these experiments we chose a parasympathetic target to test for the interesting possibility of generating compartment-specific effects, mainly due to the density of parasympathetic projections within murine islets ^4^, their release of the potent vasodilator VIP ^32^, and their clear capacity to modulate islet secretion ^1,2,32^. We did ultimately find a distinction between microvascular responses in the acinar and islet compartments, but at the parameters tested they were surprisingly subtle: while an increase in capillary motility by multiple measures was observed in both, the effect sizes were consistently larger in the acinar space, and only the acinar response approached dilation, in agreement with much evidence describing a more complex regulatory scheme in the islet capillary plexus ^6,33^. More striking was our discovery that vagal activation eliminated all traces of fast-wave vasomotion in a lobular arteriole-venule pair, consistent with much evidence that sympathetic tonus underlies this phenomenon ^34,35^, and implicating the same in adjoining capillaries. Vasomotor oscillations are thought to subserve a variety of microvascular parameters such as flow resistance, capillary flow stoppage, and tissue oxygenation but the specific physiological roles remain highly debated ^18,35,36^, so while functional consequences of these disruptions are likely, more studies are needed before specific conclusions can be made. Altogether, we believe our current findings add to the mounting evidence that vagal projections exert enough reversible influence on pancreatic microvascular activity to warrant further exploration of the potential for pancreatic neuromodulation to be of therapeutic benefit.

For this technology to mature, further characterization of the compartment-wise microvascular and secretory effects of stimulating pancreatic nerves will be needed. Reasonable next steps for this might include: targeting more local parasympathetic and sympathetic projections into the pancreas, systematically exploring stimulation parameter space for each to selectively engage distinct fiber types, and testing the capacity to combine sympathetic and parasympathetic targeting to exert bidirectional control. We specifically used long stimulation pulse trains here for maximal effect based on electrophysiological properties measured from pancreatic neurons and the time course of extended excitatory post-synaptic potentials ^2,9,10^ and imaging studies of gastric arteriole vasodilation in response to vagal stimulation lasting tens of seconds ^37^. By cuffing the abdominal vagal trunks just anterior to the gastroesophageal junction we likely activated all major branches excluding the hepatic ^38,39^, almost certainly causing off-target actuation of non-pancreatic vasculature, which more local targeting would be expected to avoid. Finally, the microvascular effects of stimulating various pancreatic nerves must be examined specifically among the known vascular actuators in and out of the islet, particularly feeding arterioles and the so-called “insulo-acinar” portal venules ^40^. The end goal of these endeavors would be to build increasingly better models connecting peripheral neural regulation, pancreatic vascular states, and eventually secretory activity, based on sufficient functional evidence.

While this preparation can achieve highly stable upright pancreatic imaging *in vivo* in dorsal recumbency, it comes with significant challenges. After eliminating respiratory movements we were surprised to find two additional mechanical oscillations to be wary of: pancreas-specific pulsations attributable to frequency of the heart rhythm, and low-frequency peristaltic movements clearly originating from vascular linkages between the pancreas and small intestine (myoenteric reflex). The former can be reliably resolved with careful mounting, but if the latter is present it can introduce small shifts of the sample over prolonged periods (minutes); it is for this reason that precisely registering vessel segments between recordings can be difficult. Regarding imaging, linescans easily out-sample full-frame resonant scanning speeds (KHz vs Hz) but are practically limited to only a few vessels at a time, and importantly, are only feasible on optimally stable preparations if recording from the smallest pancreatic capillaries. More inroads toward prolonged total stability could likely be achieved by combining features of our preparation with creative solutions seen elsewhere, such as vacuum-sealing the coverslip to the sample ^41^ or optically compensating for sample drift online ^42^. If perfect registration could be achieved, especially sampling from known pericyte locations ^5^, many more analytical approaches from modern systems neuroscience could be employed to reach stronger conclusions about what dynamic shifts may be occurring among vessel populations in response to treatments of interest. Finally, it is worth noting that a Ti:Sapphire laser is not required to image pancreatic tissue *in vivo* and this preparation can be readily adapted to any upright microscope system given the physical restraints allow it. Basic microvessel imaging has long been possible using wide-field systems, and we routinely use epifluorescence to quickly survey for specific tissue features such as islets. It is our sincere hope that the methods and tools enclosed inspire and guide others in their own optical studies of pancreatic function *in vivo*.

## Conclusion

Herein we have demonstrated several new techniques supporting *in vivo* studies of pancreatic function: an acute surgical protocol providing broad optical access to immobilized pancreas *in situ*, and a collection of complimentary TPLSM imaging strategies and analysis tools for examining diverse microvascular dynamics from multiple compartments recorded simultaneously. Using these, we uncovered some first insights into how parasympathetic projections from the vagus nerve regulate microvascular dynamics and vasomotion in the pancreas. More broadly, these findings provide further indication of the potential for neuromodular strategies to address diseases of the pancreas via vascular control, warranting deeper investigation in the future.

## Methods

### Animal Usage and Surgical Preparation

All experiments were performed as acute procedures on adult Long-Evans rats and approved beforehand by the University of Florida IACUC. Throughout surgery and data collection, animal temperatures were maintained at 37°C with a controlled heating pad and standard physiological parameters were monitored including rectal temperature, SpO2, and pulse rate (Physiosuite, Kent Scientific), as well as expiry pCO2 (Surgivet) after tracheostomy. In addition to the procedure outlined here a full detailed protocol has been uploaded to protocols.io ([link](https://www.protocols.io/view/in-vivo-two-photon-imaging-and-parasympathetic-neu-bm3tk8nn)).

Surgeries began by inducing anesthesia with a single injection of urethane i.p. (1.4 g/kg) followed by subcutaneous injections of meloxicam (1.0 mg/kg) for analgesia and 0.9 % saline for hydration. Ventral neck access is provided by a single longitudinal incision followed by careful implantation of an arterial blood pressure transducer (Transonic) sutured into the right carotid artery, taking special care to avoid damaging the cervical vagus nerve. Afterward, a standard tracheostomy is performed and the animal is placed on mechanical ventilation (Rovent, Harvard Apparatus) maintaining 80-100 bpm and end-tidal pCO2 between 20-30 mmHg for the remainder of the procedure (this eliminates the impulse to manually breathe). After ventilation is stabilized, a laparotomy is performed via a single T-shaped incision running along the linea alba, providing wide access to the diaphragm as well as anterior abdominal cavity. A bilateral pneumothorax is carefully created via a single horizontal incision through the diaphragm to bring the pleural cavity to atmospheric pressure and eliminate respiratory movements of the thoracic wall. In experiments including stimulation, the abdominal vagal trunks are accessed by implanting the esophagus just anterior to the gastroesophageal junction with a custom-made bipolar stimulation cuff electrode made from silicone tubing and coated platinum wire (4-5 kΩ impedance). In all cases, stimulation is provided as biphasic pulse trains applied in current-clamp mode by an isolated neurostimulator (AM Systems). After cuff placement, a mineral oil/Vaseline mixture is applied for insulation and protection, and proper electrical contact to the nerves is verified by confirming stimulation (for this test: 3mA current, 800us pulses, 50 Hz, 10 second train) to consistently evoke a transient drop in systemic blood pressure.

The pancreas and spleen are then located and clear connective tissues to other structures (i.e. the gastrosplenic ligature) blunt dissected while leaving all major vascular connections intact. An aluminum imaging pedestal maintained at 37°C is carefully positioned to underlie the entire spleen and 2-3 cm of tail-end pancreas and mechanically immobilized with a custom assembly of optomechanics posts and clamps (Thorlabs). Organ and pedestal positioning is fine-tuned until complete mechanical stability of the pancreas is visually achieved without any stretching, compression, or elevation changes relative the heart, and mounting is finalized by sparsely adhering both organs in place with cyanoacrylate glue (Vetbond). All newly exteriorized abdominal surfaces (excluding the immediate area to be imaged) are covered in saline-soaked gauze and maintained moist throughout the remainder of the procedure. One intravenous injection of dextran-FITC (Sigma, 2 MD) diluted in 0.9 % saline is provided as the vascular contrast agent for fluorescent imaging.

### Imaging and Vagal Stimulation Experiments

All *in vivo* imaging was performed with a Bruker Ultima two-photon laser scanning microscope system through a slightly-underfilled Nikon 16x water-dipping objective. Imaging access is provided via a coverslip placed directly on the pancreas surface and a hydrophobic sealant is applied around the perimeter (Vaseline) to support a water column for imaging. Excitation illumination to visualize dextran-FITC was provided at 940 nm, average power < 150 mW (Insight Deepsee+, Spectra-Physics) and emission was collected via GaAsP photomultiplier through a 525/70 nm filter. Full-frame videos were collected at 1024 × 1024 resolution as 15 Hz resonant scans, and random-access linescans were collected via galvo-galvo scans at 0.8 – 1.2 KHz (optimal scan frequency within this range depends on flow speed vessel-to-vessel in order to avoid over- or under-sampling). During experiments delivering repeated stimulation trials to the abdominal vagal cuff, imaging, stimulation triggers, and recording of physiological monitors are all synchronized via the Bruker Ultima GPIO (PrairieView). Voltage recordings collected in synchrony with imaging included: arterial blood pressure (sample rate 5 KHz), capnograph pCO2 waveform and ETCO2, and neurostimulator monitor signal.

A total of 89 full-frame recordings with concurrent vagal stimulation were collected from n=2 adult rats prepared as described, 60 of which including both islet and acinar regions. Each included 5 seconds of baseline, 10 seconds of stimulation, 20-45 seconds of post-stimulation recording, and an inter-trial interval of 120 seconds. Stimulation was delivered as 10 second trains of 800 us biphasic current-clamp pulses at 50 Hz with currents of 0, 0.5, 1, 1.5, 2, and 3 mA, with order randomized between trials. Full sets including all stimulation currents in at least quadruple replicate were collected at each FOV (at least one FOV containing both islet and acinar regions per animal) before moving on. Linescan recordings were also collected from an acinar arteriole-venule pair with concurrent stimulation following a similar procedure (n=5 3mA replicates with baseline periods included in each).

### Image Processing and Data Analysis

Our processing pipeline for all imaging data collected herein consists of initial processing steps using custom ImageJ/FIJI ^43^ scripts to extract raw diameter and/or directional data (for linescan recordings), followed by subsequent processing by MATLAB functions, all of which have been made freely available on protocols.io ([link](https://www.protocols.io/view/in-vivo-two-photon-imaging-and-parasympathetic-neu-bm3tk8nn)).

Full-frame recordings are first registered using the ImageJ plugin Turboreg ^44^. Vessel diameters are measured along a user-defined freehand line directed across all microvessels perpendicular to their principle axis, which is used to compile a distance vs time line profile image (similar to Figure 4b). The whole image is thresholded, morphological filtering is applied to remove holes due to erythrocyte shadows, and the width of individual vessel columns is measured at all timepoints to extract diameter traces. When drawing lines for measurement, islet capillary beds are distinguished from acinar as a spherically oriented plexus of increased capillary density, average diameter, and tortuosity, with an overall diameter between 75-250 μm (see Figure 1C and Figure 2B for example islet-acinar boundaries).

For linescan recordings, velocity columns (regions of the scanned line sampled along the principal axis) are divided into 50 millisecond time bins and analyzed via the ImageJ plugin “Directionality Analysis” to quantify the directional preference within the input by a Fourier components method ^45^. The resulting histograms are then fit with a gaussian to extract the peak preferred orientation, which is geometrically related to the instantaneous flow velocity during each time bin. Diameter columns in linescans (line regions scanned perpendicular to vessel principal axis) are analyzed identically to the process described above starting from line profile images.

Once populations of islet and acinar diameter traces have been measured, a panel of statistical and dynamical variables are calculated per population, per recording (Figure 3E). For each vessel diameter trace, period averages are calculated and used to calculate % changes in the population mean and standard deviation during stimulation compared to baseline values. Vessels exhibiting significant responses to stimulation, defined as any with diameter values exceeding ± 4 baseline standard deviations during stimulation, are counted and reported as a proportion of the whole population. The net direction of change among *r* responsive vessels is then calculated as

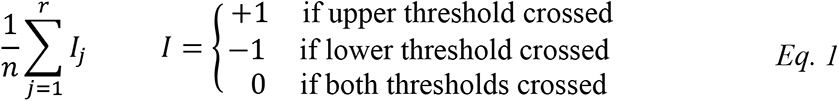

which is also normalized by the total number of vessels, *n*. Spectrograms are computed per vessel (8 second hamming window, 80% overlap), the average spectrogram is computed across all vessels, population fast-wave vasomotor power is extracted from this as the maximum value between 0.3-0.6 Hz at each timepoint, and period average powers are used to calculate % change from baseline. This window length was required to resolve the prominent 0.3-0.6 Hz oscillation band from a distinct lower frequency component also present (<0.2 Hz, visible in Figure 4C). Microvessel spectrograms are only analyzed individually in the case of Figure 4C, taken from an oscillating arteriole. The average correlation between all vessels is computed per-period and reported as a difference from baseline. Finally, histograms of diameter distributions are computed per period and the Euclidean distance, Kolmogorov-Smirnov statistic, and Kullback-Leibler divergence are computed between the stimulation period and baseline period diameter distributions.

For linescan velocity traces, period average values and % change from baseline were calculated per vessel as described above. Spectrograms were calculated from velocity data (7 second hamming window, 85% overlap) and used to measure period average vasomotor band power (0.3-0.6 Hz) per vessel.

From physiological data recorded in synchrony with imaging and stimulation, trial period averages and % changes from baseline were calculated for pulse rate (time between peaks in ABP), pulse pressure (systolic – diastolic pressure difference from ABP), and end-tidal pCO2. Raw ABP was converted to mean ABP via moving average, and the parameters of the mean ABP response following stimulation were measured by fitting the stimulation and post-stimulation periods each with sinusoids of the form

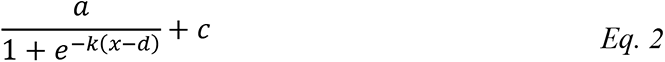

Where *c* is constrained to the end values of the preceding period and *x* is time relative to period start, thus *a* and *d* model the amplitude and time constant of the response, respectively (example fits are depicted in Supplemental Figure 3). Fitted parameters *a* and *d* during the stimulation period were output for the ABP response amplitude and ABP response delay, respectively.

### Statistical Analysis

All statistical analyses were performed in either Graphpad Prism or MATLAB using per-recording output variables as individual samples unless otherwise stated. Comparisons between >2 groups were performed by one-way Kruskal-Wallis test with reported p-values from follow-up multiple comparisons to 0 mA control group with Dunn’s correction. Modeling results in Figure 3F and Supplemental Figure 4 were generated via multiple linear regression, fitting either the stimulation amplitude or the observed ABP response amplitude per recording with the following output variables (described above) as predictors: mean diameter % change, diameter SD % change, proportion responsive vessels, net responsive direction, mean vasomotor power change, mean diameter correlation change, and all three diameter distribution change metrics. All regression p-values were obtained via standard ANOVA versus a constant model. Other tests for differences used include: Mann-Whitney p-value in Figure 3D right, paired Wilcoxin p-values in Figure 4F, and the significant ranges in Figure 4D were obtained by multiple t-tests with Holm-Sidak correction. Differences are considered significant in all cases if p < 0.05.

## Acknowledgements

We owe our gratitude to many for helping bring this work to light. Dr. Rick Johnson is thanked for much valuable guidance and input involving surgical techniques, peripheral neuroanatomy and neurophysiology, and for many productive conversations as the project matured. Dr. Martha Campbell-Thompson is thanked for orchestrating the great collaborative effort and diverse team from which this work was born, and for expert advice involving the pancreas. Vicki Dugan is thanked for patient surgical training of multiple trainees. Dr. Kristina Grove’s helpful guidance with the more technical aspects of rodent surgery, especially ventilation, were key to succeeding with the pneumothorax and achieving stability goals. Dr. Deo Singh is thanked for his participation and input during our first steps taken developing the preparation.

## Author Contributions

KO conceived the study. JC, NS, and RC developed the surgical protocol and contributed to data collection. JC and KO designed the stimulation experiments and analysis. JC implemented and carried out the analysis. AS and NS helped with raw data processing. JS and KO wrote the manuscript.

## Footnotes

a Likely extendable with the use of red fluorophore dextran conjugates and appropriate excitation source.

b Oscillating vessels were not encountered in our studies until isoflurane was replaced with urethane as the main anesthetic

