## Supplemental Figures for "*In vivo* two-photon imaging and parasympathetic neuromodulation of pancreatic microvascular dynamics in rats"

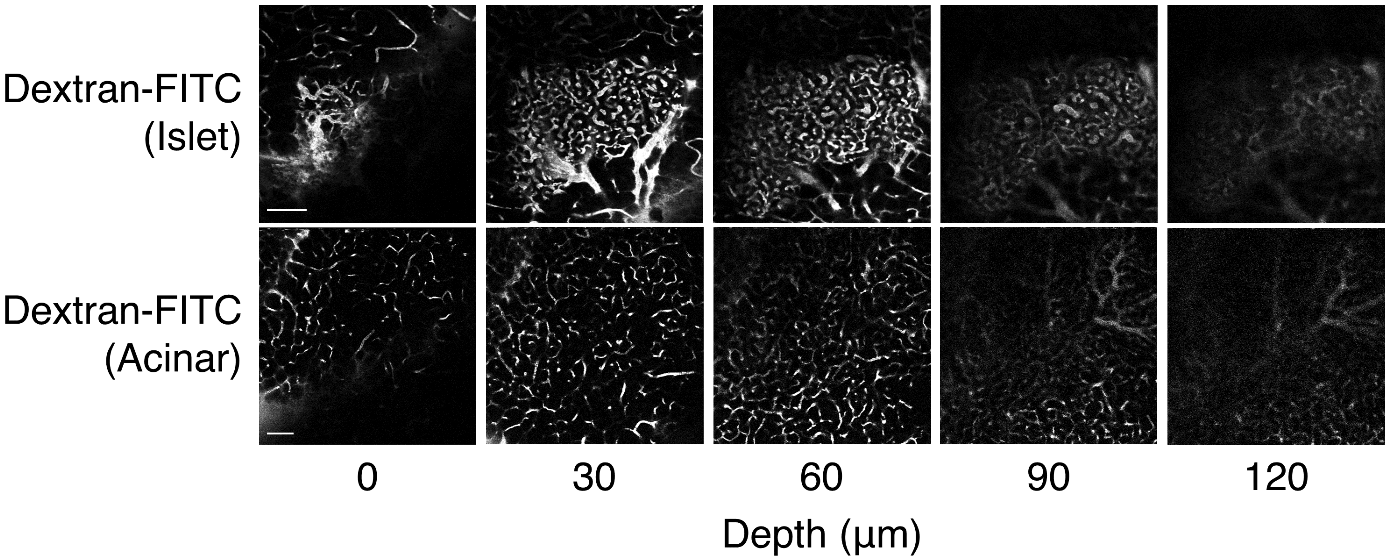


**Supplemental Figure 1** Pancreatic imaging depth achieved using TPLSM and a green fluorophore (Dextran-FITC). Top row, images of islet and peri-islet acinar capillaries taken at various depths below pancreatic surface. Scattering for individual capillaries becomes too significant for reliable measurement around 100 µm depth. Bottom row, similar procedure in an acinar-only FOV yielding similar results. Scalebars, 100 µm.

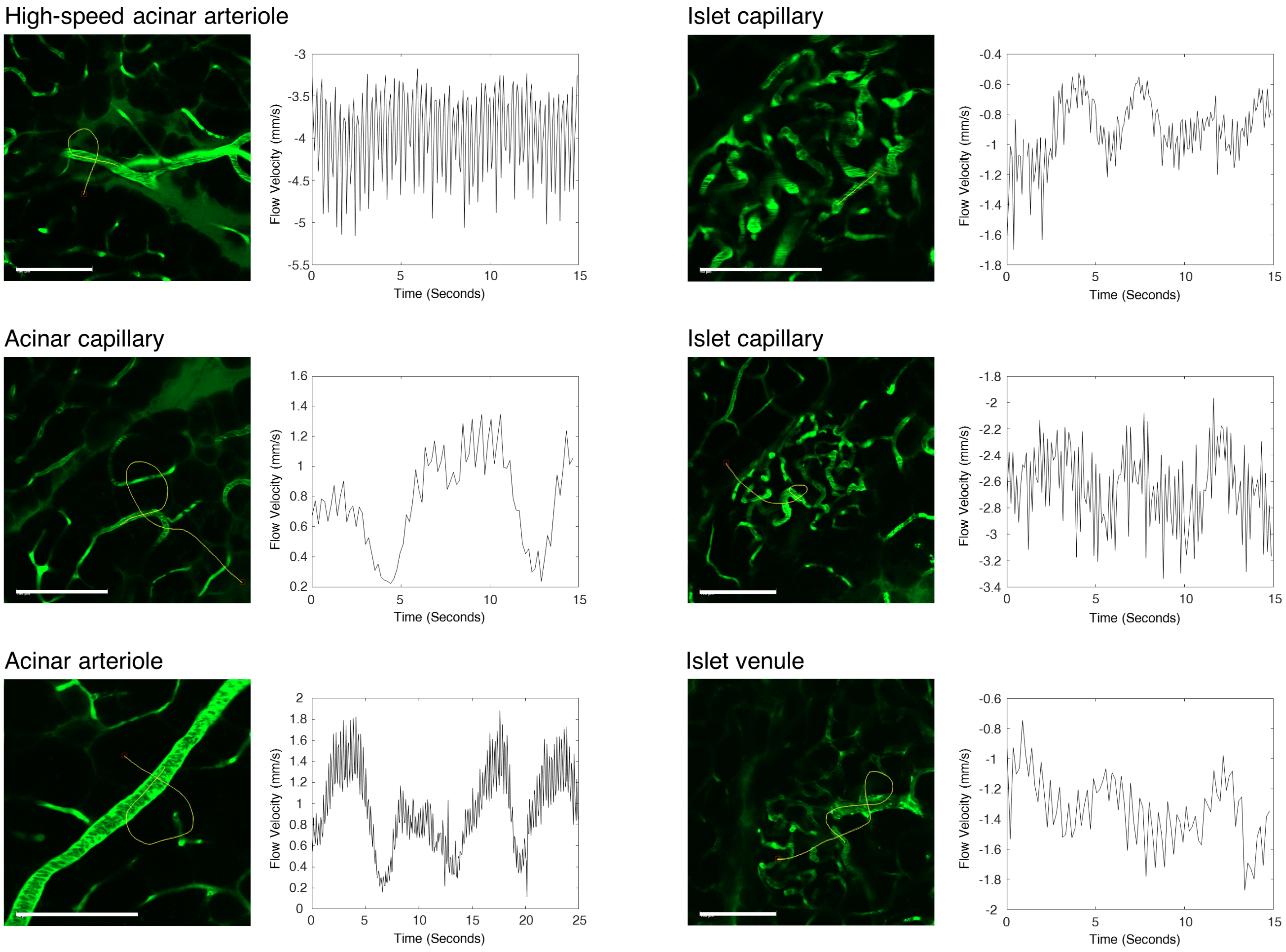


**Supplemental Figure 2** Additional flow velocity examples recorded from a variety of pancreatic microvessels. Note the common appearance of low-frequency oscillations. Vessels here were categorized by 1) size, 2) strength of cardiac oscillations in the velocity trace, and/or 3) strength of vasomotor oscillations, the latter two being traditionally attributed to arterioles rather than venules in this size range [18] (see Figures 2d and 4e for velocity traces from acinar venules). Large islet microvessels are easily classified as arteriole/venule due to the direction of flow inward/outward through the mantle. Scalebars, 100 µm.

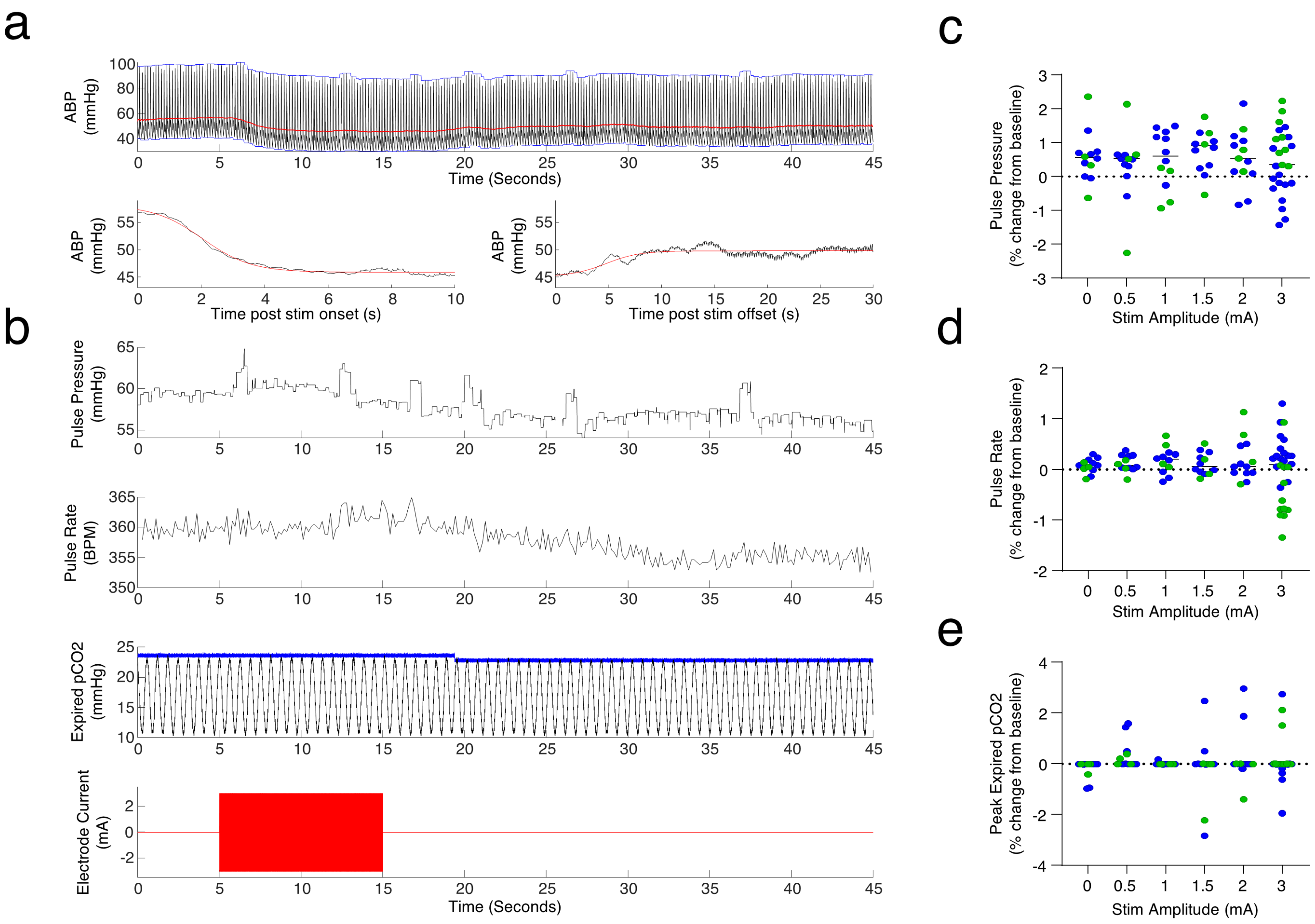


**Supplemental Figure 3** Secondary physiological measurements show no unintended cardiac or respiratory effects of abdominal vagus stimulation. (A-B) show all secondary physiological measurements performed from an example stimulation + recording experiment (3 mA) stimulation. A) Arterial blood pressure (ABP, top) recorded from a catheter transducer in the carotid artery was lowpass filtered (red line), partitioned by trial period (baseline, stimulation, post-stimulation), and sigmoids were fit to the stimulation (bottom left) and post periods (bottom right) to measure the amplitude and time course of the ABP response to stimulation (red lines are model fits). B) Other physiological measurements included pulse pressure (first row), pulse rate (second row), and peak expiry pCO2 from a capnograph in the ventilator circuit (row 3, black is capnograph waveform, blue is running maximum). Bottom row is the recorded monitor signal from the stimulation amplifier. (C-E) Vagal stimulation leads to no change in pulse pressure (C), pulse rate (D), or peak expired pCO2 (E) compared to 0 mA control trials.

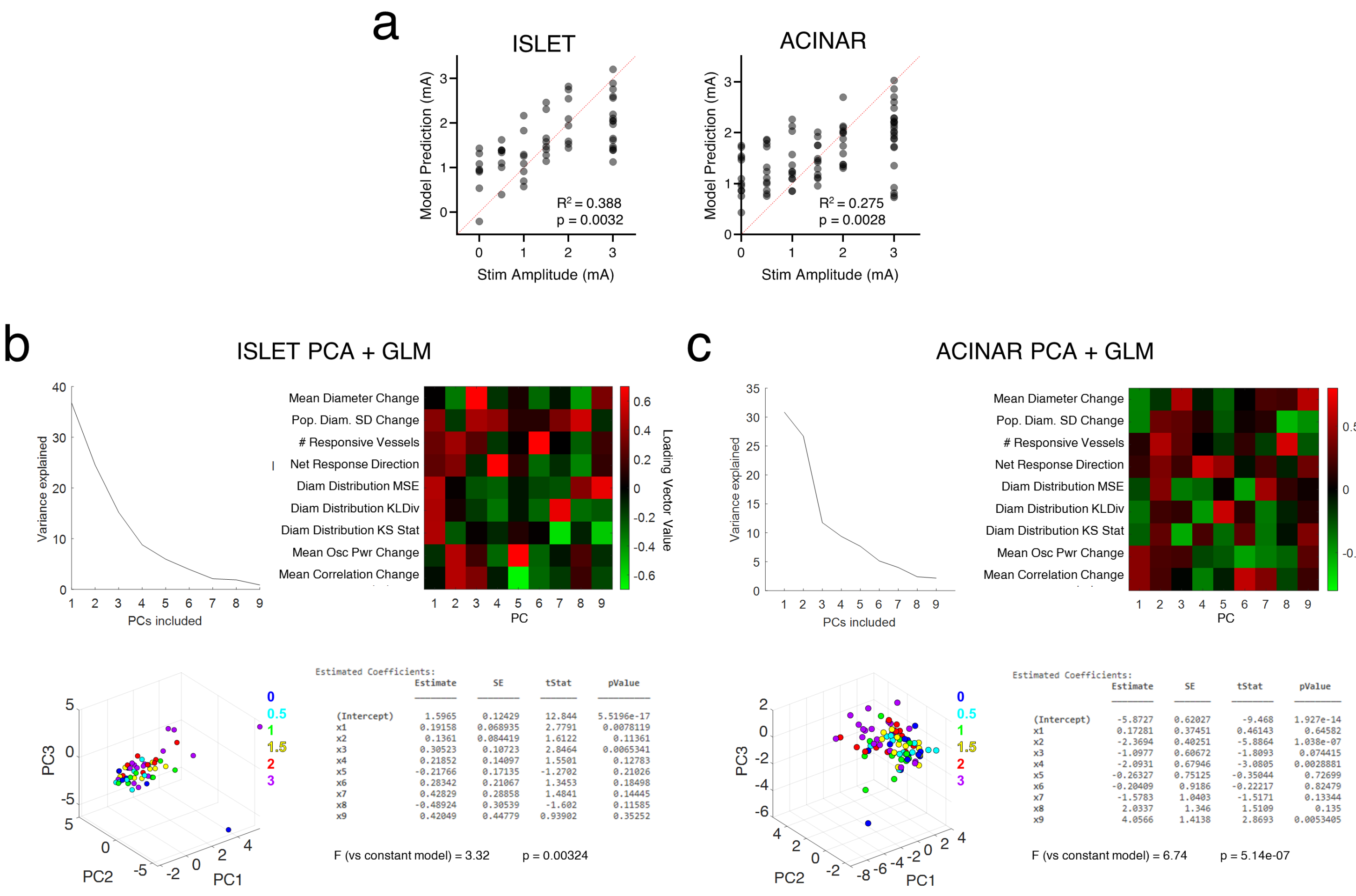


**Supplemental Figure 4** Additional modeling results and unsupervised exploration of microvascular dynamics variables from full-frame data (Fig 3). A) The microvascular dynamic variables in Fig 3E can predict the stimulation amplitude applied, using either acinar or islet microvessel data (p-values are vs a constant model). (B-C) As an additional unsupervised exploration of the data, the microvascular dynamic variables from either islet (B) or acinar (c) microvessel populations were PCA-transformed before linear fitting against ABP response amplitude, same as the procedure described in Figure 3F. For each: Top left, % variance explained for each principal component (PC). Top right, loading vectors showing the variable-wise contribution to each PC. Bottom left, scatterplot of the first three PC scores per recording colored by stimulation amplitude (legend units, mA). Bottom right, linear model coefficient fits with statistics using all PC's to predict ABP response amplitude similar to Fig 3F. Both islet and acinar PC's predict ABP response amplitude significantly better than a constant model.

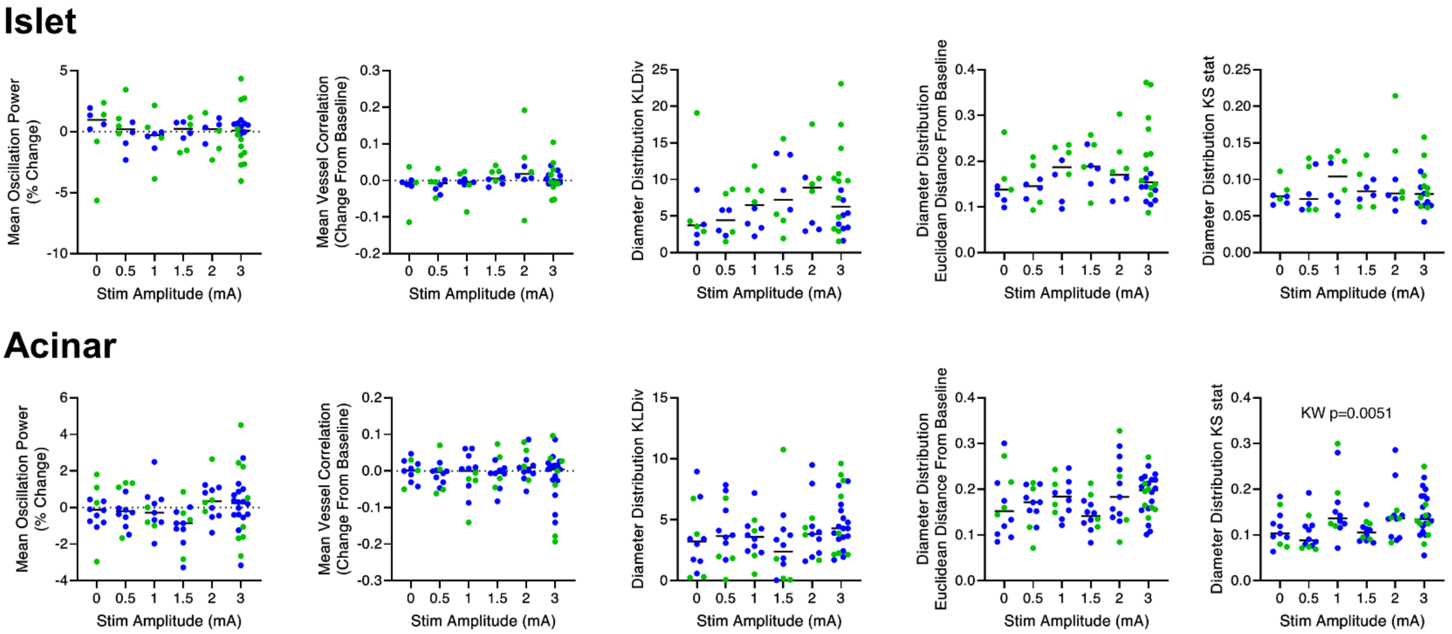


**Supplemental Figure 5** Additional microvascular population statistics plotted against stimulation amplitude for islet (top) and acinar (bottom) regions separately from full-frame data in Fig 3. All points are individual recordings colored by experiment. No significant effects of stimulation were found on the mean vasomotor oscillation power (column 1, measured within 0.3-0.6 Hz) or mean correlation between all vessels of that type (column 2), both compared against 0 mA control trials. None of three parallel metrics intended to capture changes in diameter distributions from baseline (columns 3-5 from left to right: Kullback-Leibler divergence, euclidean distance, and Kolmogorov-Smirnov statistic) were found significantly affected by stimulation compared to control trials, with the exception of acinar KS statistic (Kruskal-Wallis p=0.0051, however no multiple comparisons reached significance). Horizontal bars denote median.

**Supplemental Video 1** Raw appearance of pancreatic microvascular recordings. Left, islet and peri-islet acinar capillaries imaged in an early preparation before essential steps for stability were implemented showing strong breathing artifacts. Right, islet and peri-islet acinar capillaries imaged in a separate animal using final stability-optimized preparation. No registration was performed on either video. Scalebars, 100 μm.

**Supplemental Video 2** Vasomotor diameter oscillations and result of abdominal vagal stimulation. All scalebars are 100 μm. Stimulation current applied in the second video is 3 mA.
